## Supplemental Information for "A HaloTag-TEV genetic cassette for mechanical phenotyping of proteins from tissues"

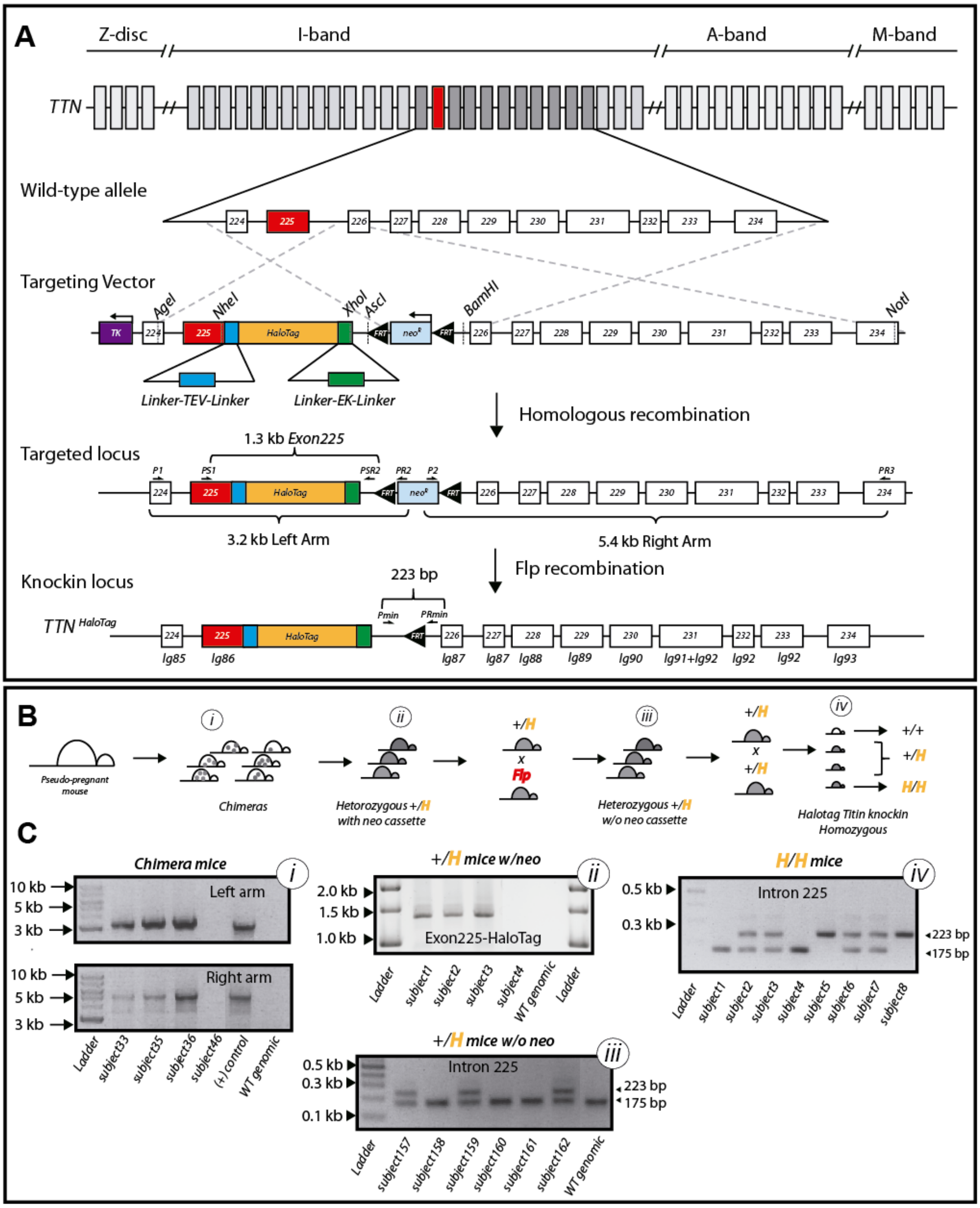


**Supplementary Figure S1. Generation of knock-in mice with the HaloTag-TEV cassette inserted in the titin gene.** (**A**) The targeting vector contains the *TTN* sequence between exons 224 and 234. The HaloTag gene is inserted downstream of exon 225, flanked by linkers including TEV and EK sites. A Neo resistance gene flanked by FRT elements was inserted in intron 225. Restriction sites and primers used during the different genetic engineering steps are shown, together with the size in base pairs (bp) of relevant fragments. (**B**) Strategy followed to get homozygous knock-in mice (H/H) from recombinant ES cells. The different experimental stages are labeled i-iv, see below. (**C**) We used PCR amplification of genomic DNA to confirm (i) the presence of the HaloTag in the TTN gene in chimera mice (primers *P1* and *PR2, left arm;* primers *P2* and *PR3*, *right arm*), (ii and iii) the presence and subsequent removal of the neo resistance by crossing with Flp mice (primers *PS1* and *PSR2,* Exon225-HaloTag*; Pmin* and *PRmin,* Intron225), and (iv) the generation of the homozygous H/H titin mouse (*Pmin* and *PRmin*). We used the vector construct and wild-type genomic DNA as controls. Sequences of primers are provided in Supplementary Table S1.

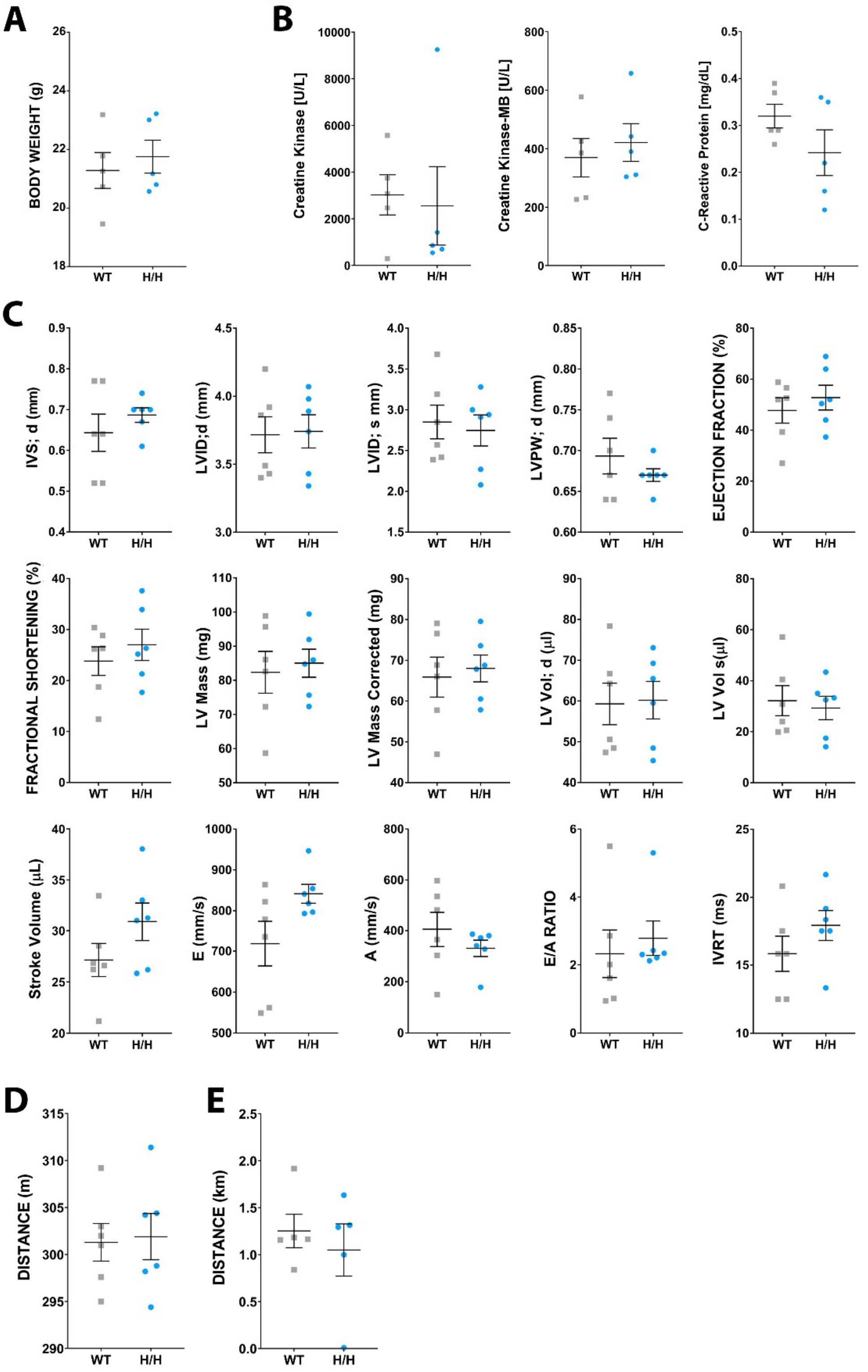


**Supplementary Figure S2. Assessment of health status of HaloTag-TEV-titin mice. (A)** Body weight of wild-type (WT) and homozygous (H/H) HaloTag-TEV-titin mice. **(B)** Serum levels of creatine kinase (marker of striated muscle damage), creatine kinase-MB (more sensitive to myocardial damage) and C-reactive protein (inflammation marker). **(C)** Cardiac function assessed by echocardiography. "s" and "d" indicate parameters obtained in systole and diastole, respectively. IVS, inter ventricular septum. LVID, left ventricular internal diameter. LVPW, left ventricular posterior wall thickness. Stroke volume, difference between endsystolic and end-diastolic volumes. E, A, early and late diastolic peak velocity waves, respectively. IVRT, isovolumic relaxation time. **(D)** Distance run by the mice during the six last training sessions. Each data point is the average value for all mice of the same genotype for a given training session. **(E)** Distance run by mice in the endurance session. In all plots, error bars represent SEM. No statistically significant difference between the groups was found for any parameter (unpaired t-test). Results in (A), (B), (D) and (E) were obtained with five 13-15-week-old female mice per group. Results in (C) were obtained with six 10-week-old male mice per group.

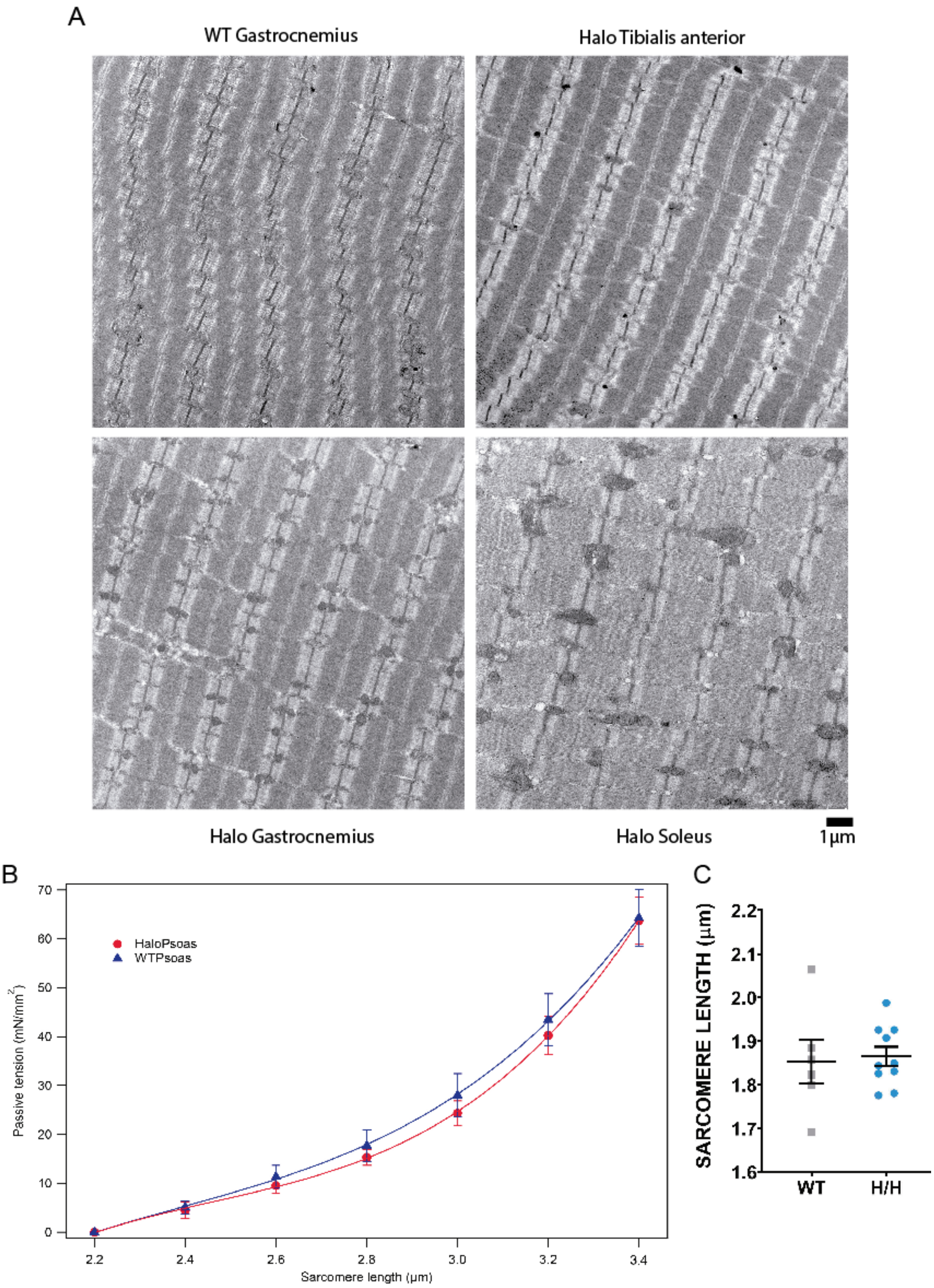


**Supplementary Figure S3. Ultrastructural and mechanical characterization of HaloTag-TEV-titin muscle fibers. (A)** The ultrastructure of HaloTag-TEV-titin muscles is equivalent to wild-type (WT). **(B)** Passive tension generated by WT and HaloTag-TEV-titin bundles of fibers isolated from psoas muscle (n = 6 for both genotypes; error bars represent SEM and the solid lines are polynomial fits). **(C)** Resting sarcomere length. n = 6 cells (WT); n = 10 cells (HaloTag-TEV-titin homozygous mice, H/H). Error bars represent SEM.

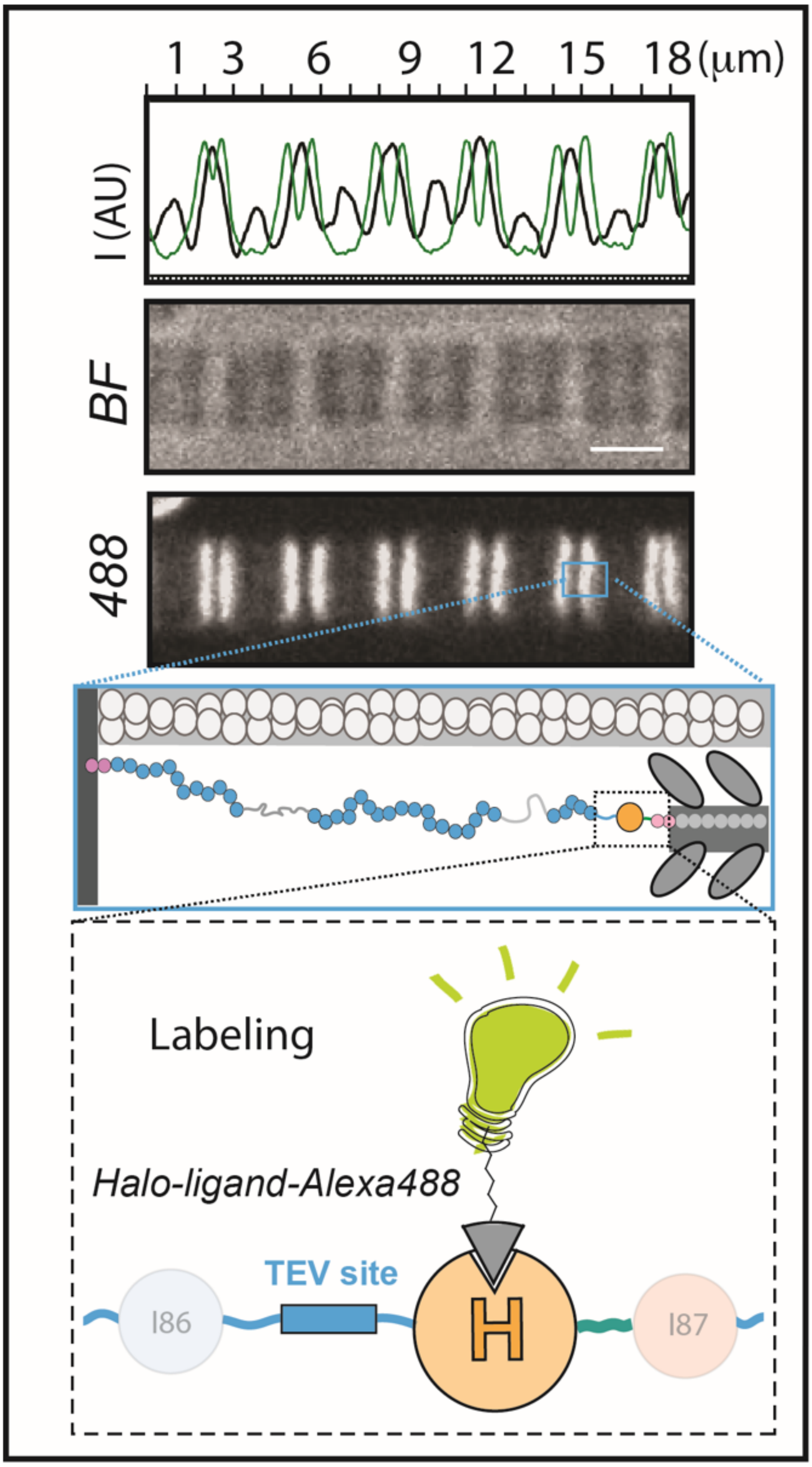


**Supplementary Figure S4. Spinning disk microscopy.** Under bright field illumination, the A- and I-band sections are easily distinguished along the myofibril as dark and clear regions, respectively (middle panel, BF). The fluorescent signal coming from Alexa488-labeled HaloTag-TEV titin appears as doublets at the I/A band interface (bottom panel, 488). The top panel shows intensity profiles (Alexa488 fluorescence in green and brightfield intensity in black). *Insets:* Cartoons showing the location of the HaloTag-labeled titin. Scale bar, 2.5 μm (also valid for Alexa488 channel).

1

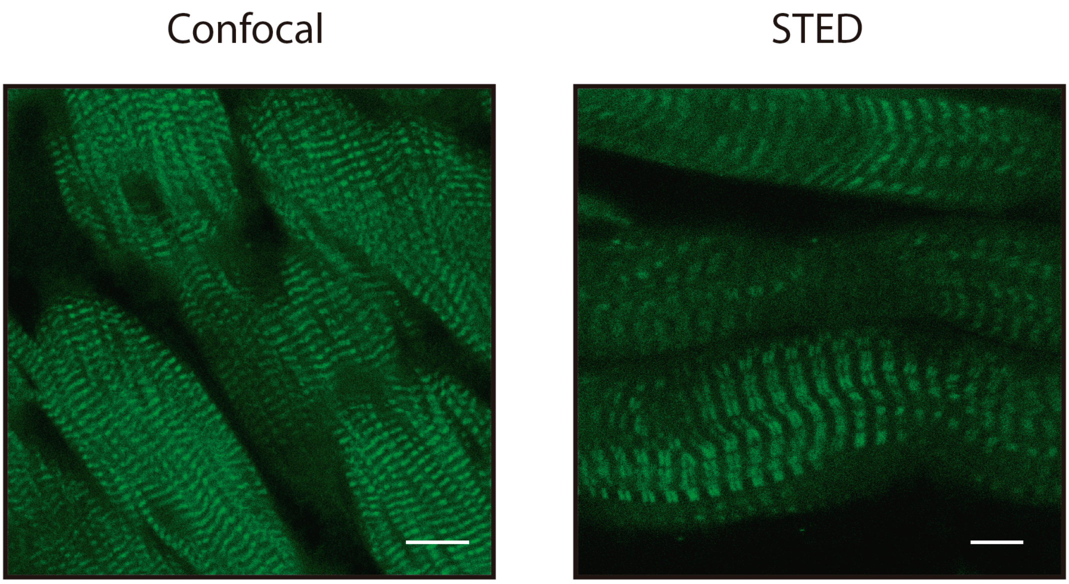


**Supplementary Figure S5. The HaloTag-TEV cassette is correctly inserted in cardiac titin.** The heart of a homozygous HaloTag-TEV titin mouse was incubated with HaloTag Oregon Green ligand, fixed and clarified. Although these samples show higher autofluorescence than skeletal preparations (**Figure 1B**), there is strong labeling in bands as expected from the location of the HaloTag insertion in titin (**Figure 1A**). Staining in doublets can also be observed, although to a lesser extent than in skeletal muscle, probably reflecting shorter I-bands in cardiac sarcomeres. Scale bars: 10 μm (Confocal) and 5 μm (STED).

1

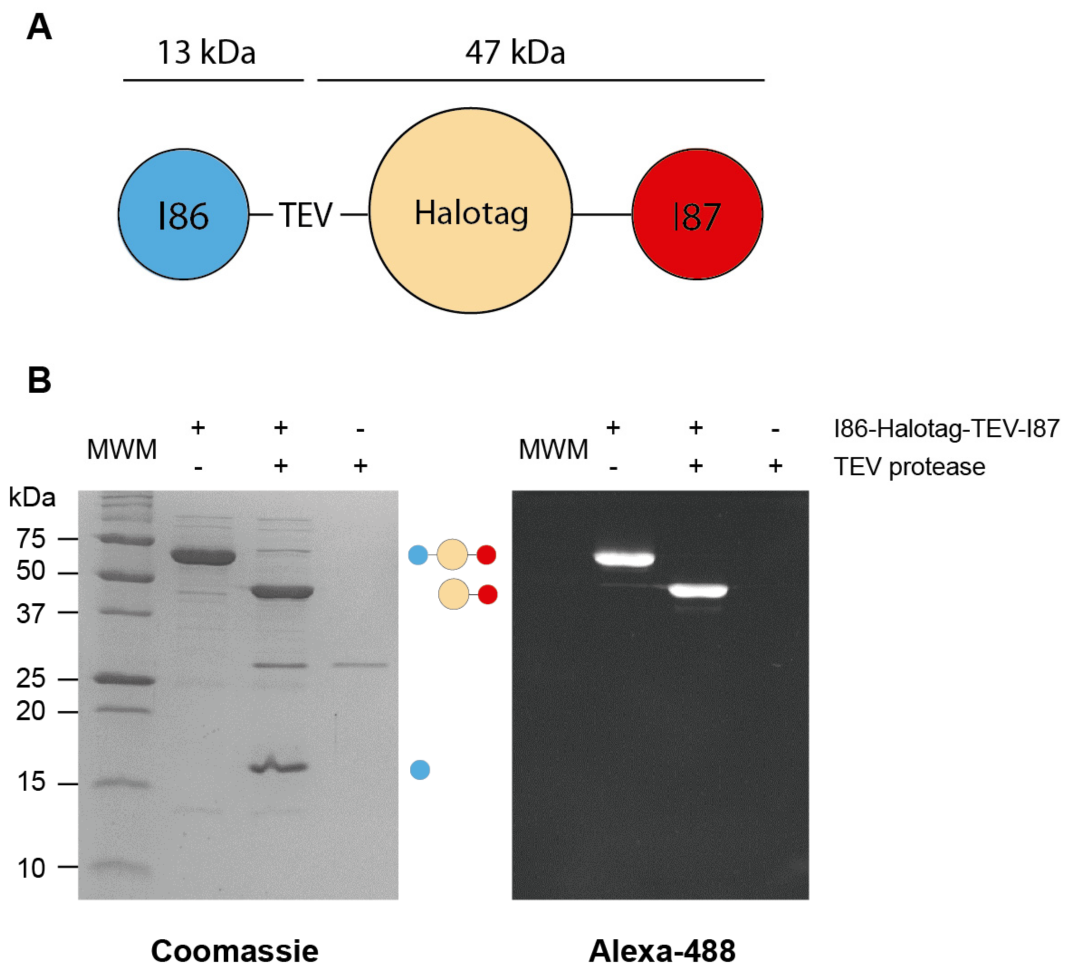


**Supplementary Figure S6. Digestion of recombinant HaloTag-TEV titin fragment. (A)** Scheme of I86-HaloTag-TEV-I87 (60 kDa) showing the size of the fragments that result from TEV digestion. **(B)** I86-HaloTag-TEV-I87 was treated or not with TEV protease (28 kDa) at 34ºC for 1 hour, and results were analyzed by 17% SDS-PAGE. Digestion resulted in the appearance of two new bands at the expected mobility (*left*, Coomassie staining). Specificity of TEV-cleavage is demonstrated by labeling with HaloTag Alexa488 ligand, which only reacts with HaloTag-containing bands (*right*, Alexa488 fluorescence).

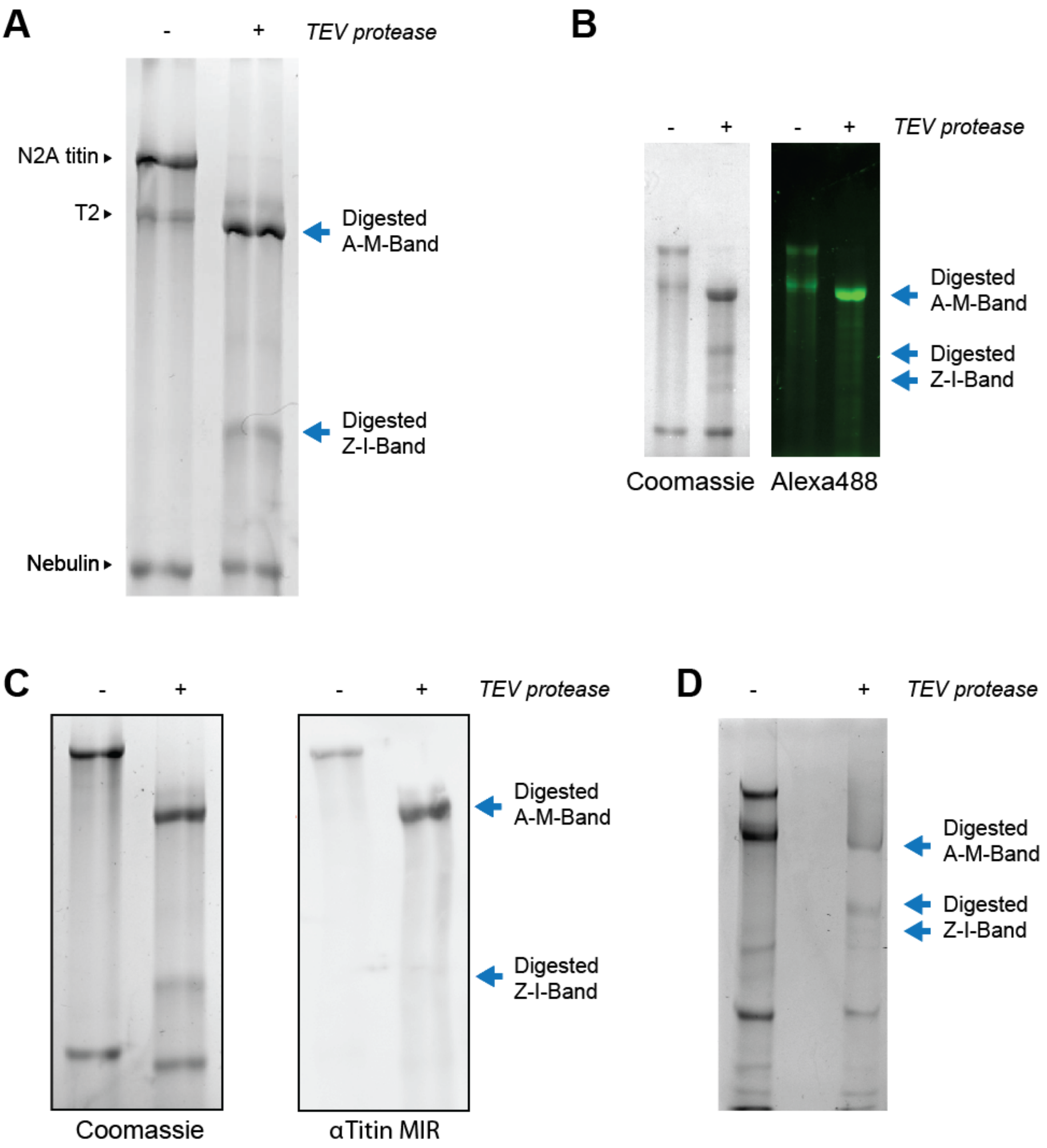


**Supplementary Figure S7. TEV digestion of skeletal muscles from homozygous HaloTag-TEV titin mice, as analyzed using 1.8% acrylamide SDS-PAGE gels. (A)** Results of TEV-digestion of isolated soleus myofibrils (Coomassie staining). **(B)** Analysis of the digestion of psoas myofibrils with TEV. Both Coomassie staining and HaloTag Alexa488 ligand are used to visualize proteins. The HaloTag-specific Alexa488 ligand only labels the digested A-M-band fragment **(C)** Equivalent results are obtained in TEV digestions of soleus myofibrils, as analyzed by western blot using the MIR antibody, which recognizes the A-band segment of titin. **(D)** The HaloTag-TEV-titin fibers used to collect the mechanical data in Figure 2C were analyzed by 1.8% SDS-PAGE to verify full digestion of titin (+ lane). A control sample with no TEV was used for reference (- lane).

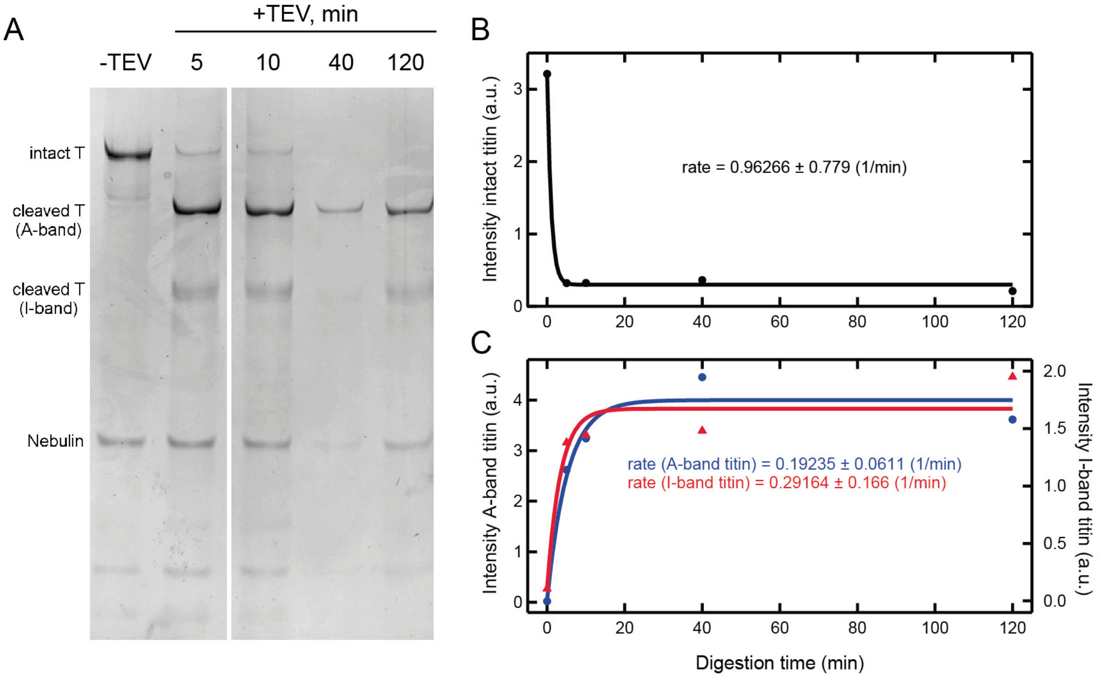


**Supplementary Figure S8. Representative kinetics of HaloTag-TEV-titin digestion by TEV protease. (A)** Psoas fibers were digested with TEV protease. Digestion was stopped at different time points by boiling aliquots in Laemmli buffer, and results were analyzed by 1.8% SDS-PAGE and Coomassie staining. Bands corresponding to nebulin, and intact and cleaved titin molecules are indicated. **(B,C)** Quantification of titin bands by densitometry. Nebulin intensity was used for normalization (black, intact titin; blue circles, A-band titin fragment; red triangles, I-band titin fragment). Solid lines are exponential fits, rate constants are indicated. These data show that at 30 min reaction time, the digestion of HaloTag-TEV titin is >99.9 % complete in this particular experiment.

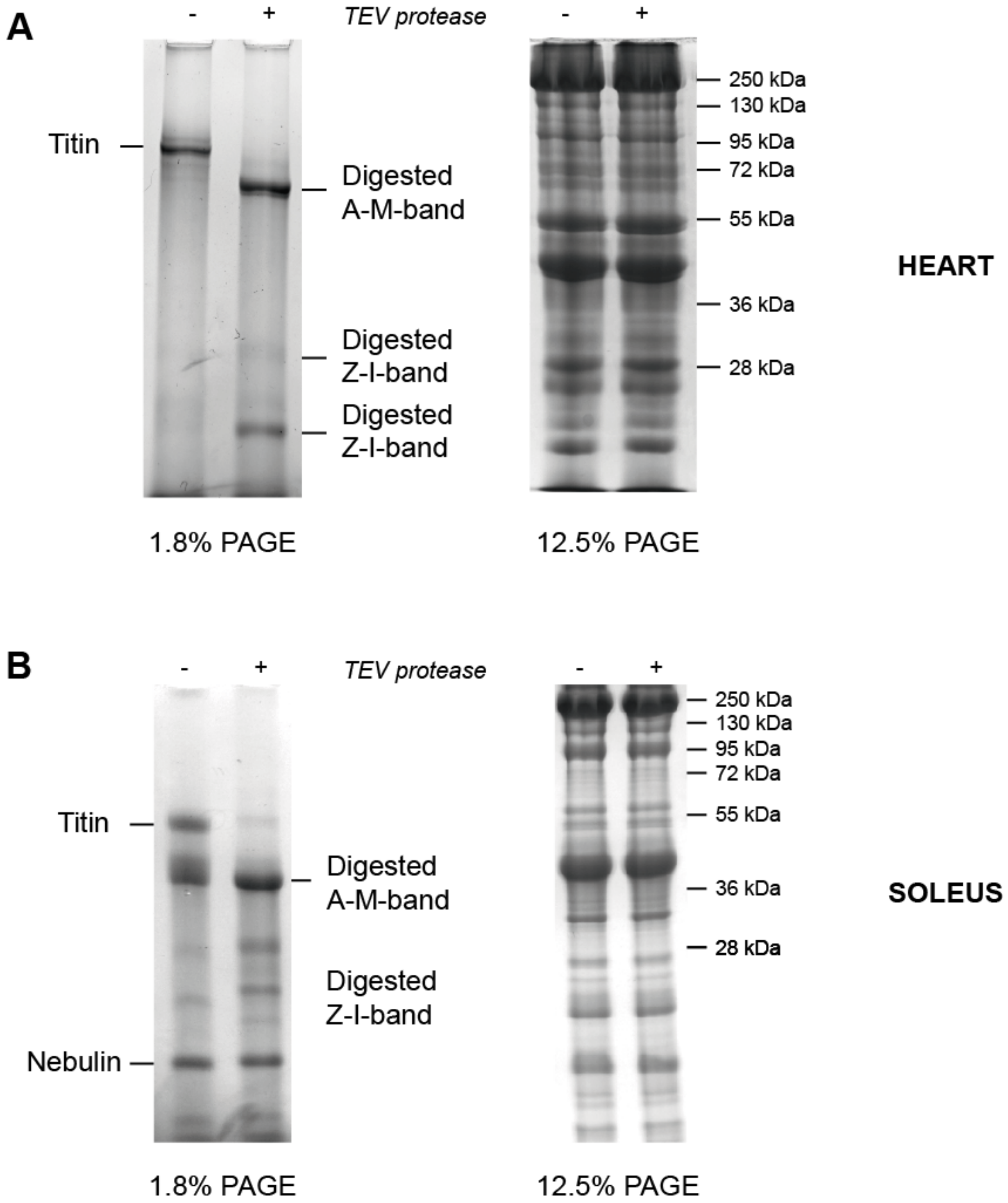


**Supplementary Figure S9. Effect of TEV treatment on the protein composition of striated muscle from homozygous HaloTag-TEV titin mice. (A)** *Left*: 1.8% SDS-PAGE shows TEV-induced specific digestion of cardiac titin. *Right*: the pattern of protein bands in 12.5% SDS-PAGE gels remains unaffected by TEV treatment. **(B)** Equivalent results are obtained with soleus samples. In this particular experiment, specific assignment of Z-I-band fragments is hindered by some non-specific degradation of titin already present in the –TEV sample.

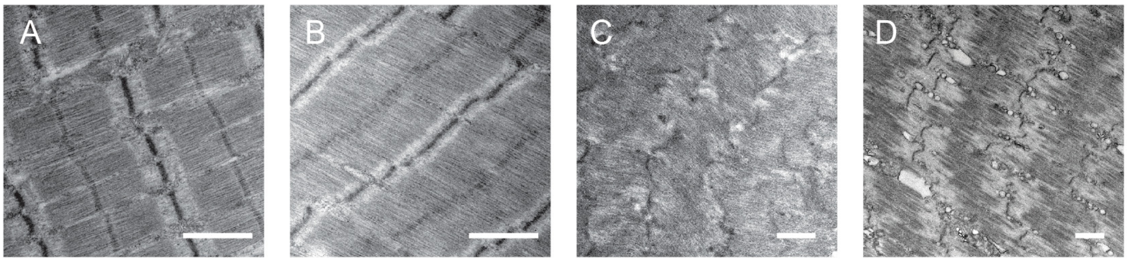


**Supplementary Figure S10. Ultrastructure of TEV-treated psoas fibers from homozygous HaloTag-TEV titin mice**. **(A)** Sample in which no TEV is addedd, fixation for EM after a stretch-release protocol. **(B)** Sample in which TEV is added, fixation in the absence of mechanical perturbation. **(C)** Sample in which TEV is added, fixation following a stretch-release cycle. **(D)** Sample in which TEV is added, fixation after holding the sample at long sarcomere length. All scale bars are 1 μm.

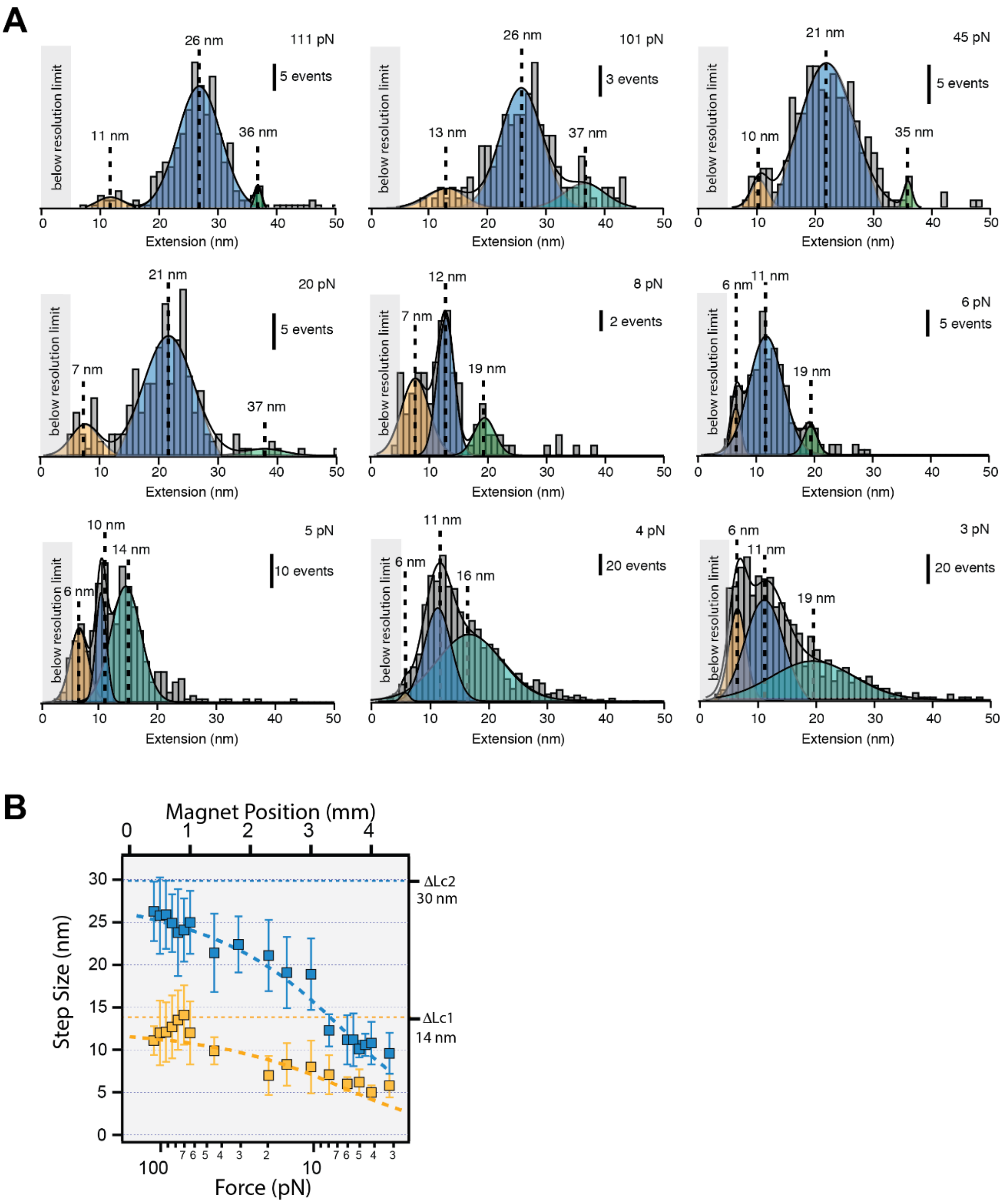


**Supplementary Figure S11. (Un)folding step sizes. (A)** Single titin molecules extracted from gastrocnemius muscles were pulled at different forces between 3 and 111 pN, and the size of the unfolding and refolding events was measured. We found three main populations of step sizes (solid lines are Gaussian fits to the data). **(B)** Force-dependency of the step sizes for the two populations corresponding to single-domain unfolding events, and fits to the worm like chain model of polymer elasticity (dashed lines). We obtain $\Delta L_C = 30 \pm 1$ nm (blue) and $\Delta L_C = 14 \pm 1$ nm (yellow).

**Supplementary Text S1. Sequence of the targeting vector to introduce HaloTag-TEV in titin.** Exons and Introns are shown in grey and white background, respectively.

Exon224—Exon225-**TEV**-HaloTag-**EK**—NeoFRT—Exon226—Exon227—Exon228—
Exon229—Exon230—Exon231—Exon232—Exon233—Exon234

**Exon224**

```
GTGGTGACACGTTCAGAAGGAAGAGTTCACACGCTCACCCTGAGGGATGTGAAGCTAGAA
GATGCTGGCGAAGTCCAACTAACTGCAAAGGATTTCAAAACTCAGGCCAATCTCTTTGTG
AAAG
gtaattagaaaacttattcctaaatacacacaatgaagacaatcacaacctctttatttt
tgtgggagagtaccacagattattaaaatcaagtttccaaaagtctaattttacatttt
ggcataacaaaatgcagggtttttttgtttttgtttttgttttaatctataagcagtg
gtagtacatgcctttaaacccagcactcaggaggcagaggcaggtggatctctgtgagtt
caaggtcagcctggtctacagagtgagttccaggacatgcagggccacacatagaaactc
tctcagaaaagaaagacaaaggatggattaaaactaacatgggttttatacttacatat
tctttctgtctctgttttattattgtatttttactagttctaagtacaactgctaaaat
aatgattaccagtccatgttatgaatcattttagccattatatcctttatgcttcatat
taattttcttgtaagctgcaatcagcagaggattgccacaagtttgaaaccagcctgatc
tacatagtgggtttccaggatagctaggaaaggggaaaaaacaaacaaacaaaaaacat
agaacagtggtttattttcctctgtatatataaatcaatgacaaacatcattgtggaagg
aaaacatctaagtgaaaaatagctttacaaacaagcaaactgagacagaagcagccccgg
agatgccagagggaatcagagaaagttctggaactcttggagaagggaaacagctaataa
gtatgattctttcag
```

**Exon225**

```
AACCCCCGGTTGAGTTCACTAAGCCTCTTGAGGACCAGACGGTCGAAGAGGAGGCCACTG
CAGTACTGGAGTGTGAAGTATCCAGAGAAAATGCCAAAGTGAAATGGTTCAAAAATGGGA
CAGAAATCCTCAAAAGCAAGAAGTATGAAATCGTTGCTGATGGCAGGGTCAGGAAGCTCA
TTATTCATGGTTGTACCCCAGAGGATATCAAAACGTACACTTGTGATGCTAAAGATTTTA
AGACCTCCTGTAACCTGAATGTTGTTC
```

**HaloTag**

TGGCTAGCGACAACACCACACCTGAG**GAGGACCTGTACTTCCAGAGC**GACAACACCACACCCGAGGCC

```
GAAATCGGAACAGGCTTCCCTTTCGACCCCCATTATGTGGAAGTGCTGGGCGAGAGGATG
CACTACGTGGATGTTGGACCCAGGGATGGCACCCCTGTGCTGTTCCTGCATGGCAACCCC
ACCAGCAGCTACGTGTGGAGGAACATCATCCCCCATGTTGCTCCTACACATAGATGCATC
GCTCCAGATCTGATTGGAATGGGAAAGAGCGATAAACCTGATCTGGGATATTTTTTCGAT
GACCATGTGAGATTTATGGATGCTTTCATTGAAGCTCTGGGACTGGAAGAAGTGGTGCTG
GTGATTCATGATTGGGGAAGCGCTCTGGGATTTCATTGGGCTAAAAGAAATCCTGAAAGA
GTGAAAGGAATTGCTTTTATGGAATTCATCAGGCCTATCCCTACCTGGGACGAATGGCCA
GAATTCGCCAGGGAGACCTTCCAGGCCTTCCGGACAACAGACGTGGGCAGAAAGCTGATC
ATCGATCAGAACGTGTTCATCGAGGGAACCCTGCCCATGGGAGTGGTCAGGCCCCTGACC
GAGGTGGAGATGGACCACTATAGGGAGCCCTTCCTGAACCCTGTTGACCGGGAGCCTCTG
TGGAGGTTCCCTAACGAGCTGCCCATCGCCGGAGAGCCCGCCAACATCGTGGCCCTGGTG
GAGGAGTACATGGATTGGCTGCACCAGAGCCCTGTGCCCAAGCTGCTGTTCTGGGGCACA
CCCGGCGTGCTGATCCCCCCTGCCGAAGCCGCTAGACTGGCTAAGAGCCTGCCCAACTGC
AAAGCTGTGGACATCGGCCCTGGACTGAATCTGCTGCAGGAAGACAACCCCGACCTGATC
GGCTCTGAGATCGCCAGGTGGCTGAGCACCCTGGAGATCAGCGGCGACAACACCACACCC
```

GAG**GACGACGACGACAAG**GACAACACCACCCCCGAGACTCGAG

```
gtaagtattcctccacaggacttggcatttgcaagtcattgtagccaaaacaaccaacac
atgtaatgcatgcccctcttggcaactcacagtatggatgctgacctaacctgatacttc
acacgtttctttacaaacatccttgtgctctgtctggccttggaaagacaccaaccact
ttttactacaccgctgcgctctttgtctgtatgtggttacacccactgtgtgactatcct
gctagagatgaaaacaaaatgagtcacaaggagaagtacagaaccctgattccacttgga
agataagtttcaggatcggaaaatatctaaaaaaataagaatgaaaagttggataggaac
atacttgagaacaactgggttatagtaacgaccaatcacataatggaactgagagtaagg
gtggacaagaagctgagtaagcccaggatctttgagtcctgattctttgaatgaagaact
ctgtcgtttttactatcacttctttccctagtggagagatgcccctctctcagtatgcc
aactgctacctagctaatttgctttgctttttttctttgtccaggaaaacatttttttcgt
ggtccattttgtgcagcgtttgtttggttctgttgcttctcatctgttgacaactaaaa
cctagtggctttaactgtgcttctggcttcttgtactctgtcactccttccatgcttggg
```

cgcctctactgtatcatcctgtagagtagccccttgggatttcttccccttgtcgtctac
ttcctcttggagaaggctcagccgtggtggcttatcttctagccttccagctagcattgg

**NeoFRT**

GAAGTTCCTATACTTTCTAGAGAATAGGAACTTCGGAATAGGAACTTCAGTGGCTATGGCAGGGCTTGCC
GCCCCGACGTTGGCTGCGAGCCCTGGGCCTTCACCCGAACTTGGGGGGTGGGGTGGGGAAAAGGAAGAAA
CGCGGGCGTATTGGCCCCAATGGGGTCTCGGTGGGGTATCGACAGAGTGCCAGCCCTGGGACCGAACCCC
GCGTTTATGAACAAACGACCCAACACCCGTGCGTTTTATTCTGTCTTTTTATTGCCGTCATAGCGCGGGT
TCCTTCCGGTATTGTCTCCTTCCGTGTTTCAGTTAGCCTCCCCCATCTCCCGTCAGAAGAACTCGTCAAG
AAGGCGATAGAAGGCGATGCGCTGCGAATCGGGAGCGGCGATACCGTAAAGCACGAGGAAGCGGTCAGCC
CATTCGCCGCCAAGCTCTTCAGCAATATCACGGGTAGCCAACGCTATGTCCTGATAGCGGTCCGCCACAC
CCAGCCGGCCACAGTCGATGAATCCAGAAAAGCGGCCATTTTCCACCATGATATTCGGCAAGCAGGCATC
GCCATGGGTCACGACGAGATCCTCGCCGTCGGGCATGCGCGCCTTGAGCCTGGCGAACAGTTCGGCTGGC
GCGAGCCCCTGATGCTCTTCGTCCAGATCATCCTGATCGACAAGACCGGCTTCCATCCGAGTACGTGCTC
GCTCGATGCGATGTTTCGCTTGGTGGTCGAATGGGCAGGTAGCCGGATCAAGCGTATGCAGCCGCCGCAT
TGCATCAGCCATGATGGATACTTTCTCGGCAGGAGCAAGGTGAGATGACAGGAGATCCTGCCCCGGCACT
TCGCCCAATAGCAGCcAGTCCCTTCCCGCTTCAGTGACAACGTCGAGCACAGCTGCGCAAGGAACGCCCG
TCGTGGCCAGCCACGATAGCCGCGCTGCCTCGTCCTGCAGTTCATTCAGGGCACCGGACAGGTCGGTCTT
GACAAAAAGAACCGGGCGCCCCTGCGCTGACAGCCGGAACACGGCGGCATCAGAGCAGCCGATTGTCTGT
TGTGCCCAGTCATAGCCGAATAGCCTCTCCACCCAAGCGGCCGGAGAACCTGCGTGCAATCCATCTTGTT
CAATGGCCGATCCCATATTGGCTGCAGGGTCGCTCGGTGTTCGAGGCCACACGCGTCACCTTAATATGCG
AAGTGGACCTGGGACCGCGCCGCCCCGACTGCATCTGCGTGTTCGAATTCGCCAATGACAAGACGCTGGG
CGGGGTTTGCTCGACATTGGGTGGAAACATTCCAGGCCTGGGTGGAGAGGCTTTTTGCTTCCTCTTGCAA
AACCACACTGCTCGACATTGGGTGGAAACATTCCAGGCCTGGGTGGAGAGGCTTTTTGCTTCCTCTTGcA
AAACCACACTGCTCGCAATTCGAAGTTCCTATACTTTCTAGAGAATAGGAACTTCGGAATAGGAACTTC

ttcctcttggagaaggctcagccgtggtggcttatcttctagccttccagctagcattgg
tctgagagcctccatgatcaagaagggttggcttggaaaggcttcttaaaaagctctgaa
agatggaagggcgaggatgtggcttctccatcctgtacagcgccagacagatttcttgga
gatgcatgaaccaacagatgtaaaagctgaatcagtgtcctatacagaacagagtttgtc
acactagtacatgcctgacttgcatatgaagtctttaaagctgagttaaaaatgaaaagc
tggatatacatggttccaattagactcgtgcctcgtagctatacatacttcaaactatta
tctttgaattctattgggtttacaatgatgtgtctttccttactagtgttccctttgaca
aagaattatccttcaactctggtatagtaccaaaacaaagaaggcatttgcttctgtaat
cattcaaaagaactgaaatgggtcccaatttgaaagaactaagttggtagcagagtaaga
tataatcgaagagaactagaaagagaacaagagcaaagcattattcccattggaagctgc
aaaggttttcaaagtcagtggcagtttatcaagacaagcacaaacagaacatgactttga
tcctagcacccaggaggcacaggcagtttggtctctgagttcaaaaccagcctggtctac
agagtgagttccaggacagccagggctacagagaggaaactctgtcttggagaaagacaa
agaaaatcatcttttcttattgtataccaaagaaatattgggataaatttgcagaagcct
catttgaattccaacattattctcttacag

**Exon226**

CTCCTCATGTGGAATTCTTAAGACCACTCACAGACCTCCAAGTCAAAGAAAAAGAAACTG
CTCGGTTCGAGTGCGAAATTTCCAAAGAAAATGAGAAG

gtctgtaacaacataagtgttaattaatattagtaagtctgattatcagtttgctttgtt
tactattgaccaaaatgcaaggaaagaagaaactagctaagcctttctgacttccatttt
agattaaatttatgctagtattaaagacagacagatgtaattgttctctttagtgccact
cattactgtctcataaattttggactcaagatatttattattgtgctggagagatggctc
agcggttaagaacactgattgctcttccagaggccctgagttcaattcccagcctccaca
tggtggctcacaaccatctgtaatggggtctgatgccctcttctggtgtgtctgaagaga
gcaatagtatactcatatacataaaataattaaataaatctttttaaaaaaagatatttg
ttattttgagtcacatggaattgatgttttcttagattaaaaaaaaagtcaaatagtagt
aattctatgtctcagctaagtagttctggttgccacaaactattattgccaaacagactt
tctgtttacatcagtggagatggtttgtgtgatatctaaggccccttaaacacaaaact
tgcataacctcatcatttgtttttccataattggaagaatatcacagtagatcaactttg
ttcttgattttaaattttattcatatttttctatcttcctttatacatactttataca
tacatataatatttatagccctcttttaaatatactataatgtcttatgattgattttaa
aaggacccttgggatactagttcatgtcatggatactgtgtaaaggaacaacatgaaaac
tgattttaagaaatctatttgtcttaaaatatag

**Exon227**

GTTCAGTGGTTTAAAGATGGTGCTGAAATTAAAAAGGGCAAAAAGTATGACATCATTTCT
AAGGGAGCAGTACGAATTCTTGTCATCAACAAATGTCTACTGAATGATGAAGCAGAATAT

TCCTGTGAAGTGAGGACAGCAAGAACTTCCGGCATGCTGACAGTCCTAG
gtgaatgtgaaggctttcttttactaagcatactagtcaggaaacccaacttccagtttt
actgactgcatctcggttcttttctgcag

**Exon228**

AAGAAGAAGCTGTCTTCACAAAAAATCTTGCCAACCTTGAAGTTAGTGAAGGAGACACTA
TCAAACTGGTGTGTGAAGTCTCCAAGCCTGGGGCAGAAGTGATTTGGTACAAAGGGGATG
AGGAGATCATCGAAACAGGGAGATTTGAAATACTTACTGATGGAAGGAAGAGAATCTTGA
TCATTCAGAATGCGCAGCTTGAGGATGCAGGCAGCTACAACTGTCGACTCCCAAGTTCTC
GAACGGACAGCAAAGTCAAAGTACACG
gtatgaaatctcagtgaagggcattttcttcctttcatccattcttgtcgtattaagata
caaatgccgatgttttcactgtcttccaactttttctttccag

**Exon229**

AACTTGCTGCTGAGTTCATCTCGAAGCCTCAAAACCTTGAAATTCTTGAAGGAGAAAAGG
CTGAGTTTGTCTGCACTATCTCAAAGGAAAGCTTCGAAGTTCAGTGGAAGAGGGATGATC
AGACACTTGAATCTGGAGATAAATATGACATCATTGCTGATGGCAAAAAGAGAGTCCTAG
TTGTAAAGGATGCCACATTACAAGACATGGGCACTTACGTAGTCATGGTTGGGGCTGCCA
GAGCCGCAGCTCACCTGACAGTCATTG
gtaagtttgtccttgctcccctgcaggcttaaatctgtagttttgatccttcatcatact
acaaagtcttttgattgtttacag

**Exon230**

AAAAACTCAGGATCATAGTTCCTCTTAAGGACACCAAGGTGAAGGAACAACAAGAGGTTG
TCTTCAACTGCGAAGTCAATACTGAAGGTGCCAAAGCCAAATGGTTCAGAAATGAAGAAG
CCATATTTGATAGTTCAAAATACATCATTCTCCAAAAAGACCTGGTCTACACCCTCAGAA
TCAGAGATGCACGGTTAGATGACCAAGCCAACTTTAATGTGTCTTTGACCAATCACAGAG
GTGAAAATGTTAAAAGTGCAGCCAATCTAATAGTGGAAG
gtatgtgacatacatgacactagagaattttcccgccagcaatttatattatgccgacaa
ctaaattcaaccttgtttttctcgatcaccacag

**Exon231**

AGGAAGATCTTAGGATTGTTGAACCTCTTAAAGATATTGAAACAATGGAGAAGAAGTCAG
TCACATTCTGGTGCAAGGTGAATCGTCTCAATGTGACACTGAAGTGGACCAAAAATGGAG
AAGAAGTGGCTTTTGACAACCGTATATCATACCGAATTGATAAGTACAAACACTCTCTAA
TCATCAAGGACTGTGGCTTCCCAGATGAAGGTGAATACGTCGTCACTGCTGGGCAAGATA
AATCCGTGGCAGAGCTGCTCATCATAGAAGCCCCAACAGAATTCGTGGAGCACCTGGAAG
ACCAGACGGTCACAGAGTTTGATGACGCTGTCTTCTCCTGCCAGCTCTCCAGAGAGAAAG
CGAATGTAAAATGGTACAGAAATGGAAGAGAAATCAAGGAAGGCAAAAA
gtacgcaaaacgtgtctgccctccctctgttgttctgtgtgtactttgacatcagacatt
tgcctaatactcatgctacaagctaactaatcattcaatctcttctttag

**Exon232**

ATACAAGTTTGAGAAGGATGGGAGCATCCACAGGCTCATCATAAAAGACTGCAGGCTGGA
GGATGAGTGTGAATACGCTTGTGGTGTAGAGGACCGCAAGTCCCGAGCTAGACTTTTTGT
AGAAG
gttagtattggcttcaaggataattctagctgaagtgacaatcttttacaatgctaata
aaaaatacaaacacatatctgttttattttacattttctctccag

**Exon233**

AAATTCCAGTTGAGATTATCAGGCCTCCTCAAGACATTCTTGAAGCCCCTGGTGCAGACG
TTATCTTCTTGGCTGAGCTCAACAAAGATAAAGTGGAGGTCCAATGGCTTAGAAATAACA
TGATCGTCGTCCAGGGTGACAAGCACCAGATGATGAGTGAAGGAAAGATACACAGGCTAC
AGATTTGTGATATTAAGCCACGTGACCAGGGCGAATACAGATTCATTGCCAAAGATAAAG
AAGCCAGGGCTAAACTTGAATTAGCAG
gtaaattgctttcttctgcttcctggtgtcctccccatatcacaaacctctgtagatttg
agattttacaattaaggcaaaacactccttgtgaaagcatatggctcgaacctgtgctca
cgtgttagctattctactttgtctacctacaaagtggaagtggtgagcagcatctttaca
tgtcaaaccactgttattcctggcatgggcagctgctcatgacttcctagaagagttcct
cactcaactgataggcaactcctgtggctcttttaaatatgaaacctgcataggtttccc
cccaaagccccattgtacatgagagaaagctgtctgacttatcctgaactaagtatgcag
gactttatcaaaaacatattttataaaataatttcaacaagaaaagcaaaacttgtttca
ccatattctaggtcctctgcaattatttttaactaagaccagatgtgaatgatggtatgc
catcttcaccaaaccactgtttccaaacatgaatagttaggagtaaaacaataacaacaa
caacaagctattataactctccaacagaatgcactgctatgaaattactgtggttatgat
aatactgtggtttatcaaatgtatagaaaaaaaaacctatgtcatagtatgactatagta
tataactcaaacatgtttgtgtgttctctttgtcactag

**Exon234**

CTGCACCTAAAATCAAGACAGCTGATCAAGATCTCGTCGTTGATGCTGGCCAGCCTCTGA
CAATGGTGGTACCCTATGATGCCTACCCCAAAGCAGAAGCTGAATGGTTTAAAGAGAACG
AACCTCTATCTACAAAAACCGTTGACACTACGGCTGAGCAGACTTCTTTCAGAATCTCAG
AAGCCAAGAAGGACGACAAGGGGAGGTATAAAATCGTGCTTCAGAACAAGCATGGGAAAG
CAGAGGGCTTCATCAATTTACAAGTTATTG

**Supplementary Text S2. Sequence of the I86-HaloTag-TEV-I87 recombinant construct.** The QS peptide bond cleaved by TEV is indicated. Please note that DNA sequence do not match the murine DNA because it was codon-optimized for optimal protein expression in *E.coli*. The first alanine residue of I87 in the HaloTag-TEV-titin construct is actually a proline in wild-type I87.

I86-TEV site-Halotag-EK site-I87

**cDNA**:

ATGAGAGGATCGCATCACCATCACCATCACGGATCCCCTCCGGTTGAATTTACCAAACCGCTGG
AAGATCAGACCGTTGAAGAAGAAGCAACCGCAGTTCTGGAATGTGAAGTTAGCCGTGAAAATGC
CAAAGTGAAATGGTTTAAAAACGGCACCGAAATCCTGAAAAGCAAGAAATATGAAATTGTGGCC
GATGGTCGTGTGCGCAAACTGATTATTCATGGTTGTACACCGGAAGATATCAAGACCTATACCT
GTGATGCCAAAGATTTCAAAACCAGCTGCAATCTGAATGTTGTTCTGGCAAGCGATAATACCAC
TCCGGAAGAGGATCTGTATTTTCAGAGTGATAATACAACCCCTGAAGCAGAAATCGGTACTGGC
TTTCCATTCGACCCCCATTATGTGGAAGTCCTGGGCGAGCGCATGCACTACGTCGATGTTGGTC
CGCGCGATGGCACCCCTGTGCTGTTCCTGCACGGTAACCCGACCTCCTCCTACGTGTGGCGCAA
CATCATCCCGCATGTTGCACCGACCCATCGCTGCATTGCTCCAGACCTGATCGGTATGGGCAAA
TCCGACAAACCAGACCTGGGTTATTTCTTCGACGACCACGTCCGCTTCATGGATGCCTTCATCG
AAGCCCTGGGTCTGGAAGAGGTCGTCCTGGTCATTCACGACTGGGGCTCCGCTCTGGGTTTCCA
CTGGGCCAAGCGCAATCCAGAGCGCGTCAAAGGTATTGCATTTATGGAGTTCATCCGCCCTATC
CCGACCTGGGACGAATGGCCAGAATTTGCCCGCGAGACCTTCCAGGCCTTCCGCACCACCGACG
TCGGCCGCAAGCTGATCATCGATCAGAACGTTTTTATCGAGGGTACGCTGCCGATGGGTGTCGT
CCGCCCGCTGACTGAAGTCGAGATGGACCATTACCGCGAGCCGTTCCTGAATCCTGTTGACCGC
GAGCCACTGTGGCGCTTCCCAAACGAGCTGCCAATCGCCGGTGAGCCAGCGAACATCGTCGCGC
TGGTCGAAGAATACATGGACTGGCTGCACCAGTCCCCTGTCCCGAAGCTGCTGTTCTGGGGCAC
CCCAGGCGTTCTGATCCCACCGGCCGAAGCCGCTCGCCTGGCCAAAAGCCTGCCTAACTGCAAG
GCTGTGGACATCGGCCCGGGTCTGAATCTGCTGCAAGAAGACAACCCGGACCTGATCGGCAGCG
AGATCGCGCGCTGGCTGTCGACGCTCGAGATTTCCGGCGATAACACGACACCTGAAGATGATGA
TGATAAAGACAATACGACACCGGAAACACGTGCACCGCATGTGGAATTTCTGCGTCCGCTGACC
GATCTGCAGGTTAAAGAAAAAGAAACCGCACGTTTTGAATGCGAGATCAGCAAAGAAAATGAAA
AGGTGCAGTGGTTTAAAGATGGTGCCGAAATCAAAAAAGGCAAAAAATACGACATCATCTCCAA
AGGTGCCGTTCGTATTCTGGTTATTAACAAATGTCTGCTGAACGATGAAGCCGAATATAGCTGT
GAAGTTCGTACCGCACGTACCAGCGGTATGCTGACCAGATCTTAA

**Protein**:

MRGSHHHHHHGSPPVEFTKPLEDQTVEEEATAVLECEVSRENAKVKWFKNGTEILKSKKYEIVA
DGRVRKLIIHGCTPEDIKTYTCDAKDFKTSCNLNVVLASDNTTPEEDLYFQ'SDNTTPEAEIGT
GFPFDPHYVEVLGERMHYVDVGPRDGTPVLFLHGNPTSSYVWRNIIPHVAPTHRCIAPDLIGMG
KSDKPDLGYFFDDHVRFMDAFIEALGLEEVVLVIHDWGSALGFHWAKRNPERVKGIAFMEFIRP
IPTWDEWPEFARETFQAFRTTDVGRKLIIDQNVFIEGTLPMGVVRPLTEVEMDHYREPFLNPVD
REPLWRFPNELPIAGEPANIVALVEEYMDWLHQSPVPKLLFWGTPGVLIPPAEAARLAKSLPNC
KAVDIGPGLNLLQEDNPDLIGSEIARWLSTLEISGDNTTPEDDDDKDNTTPETRAPHVEFLRPL
TDLQVKEKETARFECEISKENEKVQWFKDGAEIKKGKKYDIISKGAVRILVINKCLLNDEAEYS
CEVRTARTSGMLTRS

**Supplementary Table S1. Sequence of primers used to produce and genotype HaloTag-TEV-titin mice.**

| Primer | Sequence |
|---|---|
| **P1** | 5'-ACCCTGAGGGATGTGAAGC-3' |
| **PS1** | 5'-TGCAGTACTGGAGTGTGAAGTATCC-3' |
| **PSR2** | 5'-GAAACGTGTGAAGTATCAGGTTAGG-3' |
| **PR2** | 5'-CTGGCACTCTGTCGATACCC-3' |
| **P2** | 5'-GGGTTTGCTCGACATTGG-3' |
| **PR3** | 5'-GTAAATTGATGAAGCCCTCTGC-3' |
| **Pmin** | 5'-CGTGGTGGCTTATCTTCTAGC-3' |
| **PRmin** | 5'-CTGTTGGTTCATGCATCTCC-3' |

**Supplementary Video S1. 3D reconstruction of HaloTag-TEV clarified muscle fibers.** Following specific labeling of gastrocnemius muscle with Oregon Green Halo ligand, fixation and clarification, a 250-µm-deep Z-stack was obtained using multiphoton microscopy (127 images). Individual images were intensity corrected and the software Imaris was used to produce the 3D animation.

**Supplementary Reference**

1. Li, H. *et al.* Reverse engineering of the giant muscle protein titin. *Nature* **418**, 998–1002 (2002).
